# Serine proteases are required to activate influenza D virus haemagglutinin-esterase fusion (HEF) protein

**DOI:** 10.64898/2026.04.16.717628

**Authors:** Meshach M. Maina, Junsen Zhang, Martin Mayora Neto, Kelly da Costa, Eva Bottcher-Friebertshauser, Edward Hutchinson, Maria Giovanna Marotta, Claudia Maria Trombetta, Simon D. Scott, Nigel Temperton, Janet M. Daly

**Affiliations:** One Virology, Wolfson Centre for Global Virus Research, School of Veterinary Medicine and Science, University of Nottingham, UK; MRC-University of Glasgow Centre for Virus Research, Glasgow, UK; Viral Pseudotype Unit, School of Pharmacy, University of Kent and Greenwich, UK; Institute of Virology, Philipps University, 35043 Marburg, Germany; Department of Molecular and Developmental Medicine, University of Siena, Siena, Italy

## Abstract

Influenza D virus (IDV), the most recently identified member of the *Orthomyxoviridae*, was first isolated from pigs but cattle have been identified as the reservoir host. To date, IDV has not been confirmed to cause human disease. Like the haemagglutinin (HA) of influenza A virus (IAV) and the haemagglutinin-esterase fusion (HEF) protein of influenza C virus (ICV), the IDV HEF is produced as a precursor protein (HEF0) that must be proteolytically cleaved by host cell proteases (into HEF1 and HEF2) to gain its fusion capacity. The proteases that activate IAV HA have been extensively studied, but those responsible for activation of IDV HEF were unknown. Identifying these proteases is key to understanding early virus–host interactions and host restriction. Therefore, we generated ICV and IDV pseudotyped viruses (PVs) in HEK 293T producer cells with or without co-transfection of plasmids expressing different type II serine proteases. Subsequent transduction of swine testicular (ST) cells indicated strong activation of both ICV and IDV PVs by the human airway trypsin-like protease (HAT) and its swine homologue (swAT). Furthermore, like influenza A/Puerto Rico/8/34 (H1N1) virus, addition of exogenous protease is not essential for IDV replication in MDCK II cells, most likely due to endogenous expression of matriptase.

In conclusion, our data unveil new information on host cell proteases that activate ICV and IDV HEF proteins. Importantly, the data suggest that protease specificity is not a factor in restriction of IDV replication in the human upper respiratory tract.

## Introduction

Influenza viruses belong to the family *Orthomyxoviridae* in the order *Articulavirales*, which have a segmented negative-sense RNA genome [1]. The family currently has four genera, *alphainfluenzavirus* (influenza A virus, IAV), *betainfluenzavirus* (influenza B virus, IBV), *gammainfluenzavirus* (influenza C virus, ICV) and *deltainfluenzavirus* (influenza D virus, IDV) in which viruses have eight (IAV and IBV) or seven (ICV and IDV) genome segments.

Influenza C virus, a predominantly human pathogen causing mild upper respiratory tract infections in children, was first isolated over 70 years ago [2, 3]. The virus has been reported to infect dogs, cattle and pigs, albeit rarely [4-7]. Serological evidence suggests that ICV has a global distribution with most people developing antibodies against the virus in early stages of life [8, 9].

In 2011, an influenza virus was isolated from a pig with influenza-like signs in the United States [10]. Electron microscopy, genetic and antigenic analysis revealed that this virus was neither IAV nor IBV but possibly represented a new ICV subtype [10, 11]. However, the virus presented only 50% identity with ICV, and no cross reactivity was observed between antibodies against the virus and IAV, IBV or ICV [10]. Consequently, the virus was classified as a new genus (*Deltainfluenzavirus*) of the *Orthomyxoviridae* family [12].

Influenza D virus has a global distribution [11-13] with two antigenically and phylogenetically different clades, which have been shown to frequently reassort, identified in the USA [14], and a third clade identified in Japan [15]. Although it was first isolated from pigs, it has been shown that cattle are the amplification and primary reservoir host for the virus [12, 13]. Antibodies to IDV have also been detected in small ruminants, camels, horses, cats, dogs and humans [13, 16-22]. However, studies have suggested that limited availability of 9-O-acetylated sialic acid receptors, which are widespread in the bovine respiratory tract, may restrict viral entry in these species [23, 24]. Nevertheless, aerosol challenge of four influenza-naïve horses with 6.25×10^7^ 50% tissue culture infectious doses per animal of IDV demonstrated that the virus can replicate, with detection of virus shedding from the nasopharynx and induction of neutralising antibodies, but all four animals were asymptomatic [25].

Influenza C virus and IDV both have a single major envelope surface glycoprotein, haemagglutinin esterase fusion (HEF), which initiates infection by receptor binding and mediates fusion of the viral envelope with the endosomal membrane and degrades the viral receptor in infected cells. Like many viral fusion proteins, HEF is synthesized as a cleavable precursor (HEF0) that is post-translationally cleaved into N-terminal HEF1 and C-terminal HEF2 [26, 27]. Cleavage activation of HEF is required for the conformational changes that trigger membrane fusion at acidic pH in the endosome [28]. Therefore, this post-translational process is essential for viral infectivity and replication [27]. The specific cellular proteases that are required for the activation of viral fusion proteins depends on the amino acid sequence at the cleavage site of the protein [29] and the expression of these proteases determines the ability of a cell to produce infectious virus particles [13]. HEF has a monobasic cleavage site (a tryptic cleavage site in which cleavage occurs after a single lysine or arginine residue) and is in this respect it is similar to the haemagglutinin (HA) of low pathogenicity avian IAV and of IAVs adapted to mammalian species, IBV and ICV [27]. Influenza HAs with monobasic cleavage sites are activated by type II transmembrane serine proteases including transmembrane protease serine S1 member 2 (TMPRSS2) and human airway trypsin-like protease (HAT) [30]; proteases commonly found in mammalian respiratory tissue. In contrast, the multi-basic cleavage site (MCS) found in the HA of H5 and H7 subtype highly pathogenic avian influenza (HPAI) viruses, which can be recognised by furin or furin-like serine proteases found in many cell types, is not found in any known HEF protein [13, 28].

Which proteases activate IDV HEF have yet to be identified. Recently, Sato, Hayashi [31] demonstrated that ICV was able to undergo multiple cycles of replication in Madin-Darby canine kidney (MDCK) cells stably expressing TMPRSS2 but not in cells stably expressing HAT, which led us to hypothesise that IDV might have a similar cleavage preference.

Lentiviral reporter viruses, known as pseudotyped viruses (PVs), have previously been used to determine the requirement for proteolytic activation of IAV and severe acute respiratory syndrome coronavirus (SARS-CoV) envelope proteins [32-34]. We recently reported the development and optimisation of lentiviral PVs to facilitate the study of ICV and IDV [35]. In this study, PVs were used to investigate the activation of ICV and IDV HEF proteins by type II transmembrane serine proteases and the subsequent cell tropism of IDV.

## Materials and methods

### Cell lines

Human A549 lung adenocarcinoma cells, human embryonic kidney (HEK) 293T cells, Madin-Darby bovine kidney (MDBK) cells, Madin-Darby canine kidney (MDCK) cells and swine testicular (ST) cells were grown in Dulbecco’s modified Eagle medium (DMEM; Gibco) supplemented with 2 mM L-glutamine, 1% penicillin (100 units/ml) and streptomycin (100 μg /ml) (Sigma-Aldrich), and 10% fetal bovine serum (FBS; Gibco, ThermoFisher Scientific) at 37°C in a 5% CO_2_ incubator.

### Plasmids

The custom-synthesised HEF protein open reading frame (ORF) of C/Minnesota/33/2015 was cloned into pcDNA 3.1 plasmid (pcDNA-HEF_ICV_) as previously described [35]. Similarly, the HEF ORFs of D/swine/Italy/199723-3/2015 and D/bovine/France/5920/2014 were custom-synthesised or PCR-amplified from the virus, respectively, and cloned into the pI.18 expression plasmid to produce pI.18-HEF_Italy_ and pI.18-HEF_France_ as previously described [35].

The pCAGGS mammalian expression plasmid coding for HAT (pCAGGS-HAT), TMPRSS2 (pCAGGS-TMPRSS2), swine airway trypsin-like protease (pCAGGS-swAT) and swine transmembrane protease, serine S1 member 2 (pCAGGS-swTMPRSS2) were generated as previously described [30, 36]. The human matriptase sequence (TMPRSS14; NP_068813.1) with human codon optimisation and a Kozak sequence was synthesised by GenScript and cloned into pcDNA3.1(+) and kindly provided by Dr T. Peacock (The Pirbright Institute).

### Production of pseudotyped viruses

To produce PVs, the pCSFLW lentiviral vector coding for the reporter (firefly luciferase), psPAX2 coding for the HIV gag/pol and the relevant envelope protein expression plasmid (pcDNA-HEF_ICV_, pI.18-HEF_Italy_, pI.18-HEF_France_) were co-transfected using Lipofectamine 2000 (Invitrogen), according to the manufacturer’s instructions, into HEK293T cells in a 6-well plate. A plasmid expressing the vesicular stomatitis virus glycoprotein (pCAGGS-VSV-G) and an expression plasmid with no insert (Δenvelope) were used as positive and negative controls, respectively. To determine the activation efficiency of different proteases, pCAGGS-HAT, pCAGGS-TMPRSS2, pCAGGS-swAT, pCAGGS-swTMPRSS2 or pcDNA3.1-matriptase were co-transfected with the core, reporter and envelope plasmids. Alternatively, 1 μg/ml of TPCK-treated trypsin was added immediately after transfection. The PVs were harvested 48 hours post transfection, passed through 0.45 µm cellulose acetate filters (Minisart) and stored at -80°C.

### Transduction of target cells with pseudotyped viruses

Transduction efficiency of PVs was determined by seeding ST cells in 96-well white plates to be 70–85% confluent after overnight incubation. Subsequently, fresh media was added and the PVs were serially diluted (two-fold dilution in eight steps) in the plate and then incubated for 48 hours before adding Promega Steady-Glo and measuring the relative light unit (RLU) values with a GloMax Multi detection system luminometer (Promega). The ‘transduction titre’ was calculated as the mean of the relative light units per ml (RLU/ml) calculated for each dilution of PV. The HEF protein plasmid was omitted to produce a ‘Δenv’ control and ‘no protease’ controls were included.

### Viruses

The H1N1 subtype IAV A/Puerto Rico/8/34 (A/PR8) and D/bovine/France/5920/2014 were cultivated in MDCK cells. Replication curves were obtained by inoculation of A549, MDBK or MDCK cells with a multiplicity of infection of 0.001 with 0.5 or 1 μg/ml of TPCK-treated trypsin or with no trypsin added to the culture medium. Supernatants were harvested at 6, 24, 48 and 72 96 h post-infection and plaque assays performed on MDCK cells as described previously [36] to obtain plaque forming units per ml (PFU/ml). For the plaque assays with IDV immunocytochemical staining was used to visualise plaques. A sheep polyclonal antibody against IDV nucleoprotein (available from www.influenza.bio) was used at 1/500 dilution as the primary antibody with Alexa Fluor™ 568 donkey anti-sheep secondary (Thermo Fisher) at 1/1000 dilution as the secondary antibody. Nuclei were counterstained with DAPI and plaques visualised with a Celigo imaging cytometer (Nexcelom).

### Sequence analysis

Codon-based analysis of viral cleavage sites was performed by downloading the full-length HEF sequences of D/swine/Italy/2015 (ALE66296.1) and D/bovine/France/2014 (AUO38022.1) from the National Centre for Biotechnology Information Database (NCBI) and aligning them using the MUSCLE algorithm [37] in Molecular Evolutionary Genetics Analysis (MEGA) [38]. The protease cleavage site was then identified by comparison with the ICV HEF cleavage site sequence identified in [35] by alignment of full-length nucleotide sequence data from 352 strains between 1947 and 2018.

## Results

The full-length HEF sequences of the IDV isolates used to generate PVs (D/swine/Italy/2015 and D/bovine/France/2014) were identified by comparison with the published cleavage site sequence for ICV HEF [35]. The aligned amino acid sequences of the HEF protease cleavage site of the IDV viruses were identical (Table 1), and the three amino acids downstream of the cleavage site were identical in ICV and IDV. Apart from the arginine residue immediately before the cleavage site, present in IAV, IBV, ICV and IDV, there was limited conservation of the sequence upstream of the cleavage site.

**Table 1.** Comparison of amino acid residues either side of the protease cleavage site (P0) in influenza viruses A, B, C (taken from [31]) and the HEF cleavage site sequences of D/swine/Italy/2015 (ALE66296.1) and D/bovine/France/2014 (AUO38022.1), which were used to generate the influenza D virus pseudotyped viruses used in this study.

| Virus<br>sequence | Residue at position: |  |  |  |  |  |  |  |  |  |  |
| --- | --- | --- | --- | --- | --- | --- | --- | --- | --- | --- | --- |
|  | P-7 | P-6 | P-5 | P-4 | P-3 | P-2 | P-1 | P0 | P+1 | P+2 | P+3 |
| IAV H1 | I/V | P | S | I | Q | S | **R** | ✂ | G | L | F |
| IAV H3 | I/V | P | E | K | Q | T | **R** | ✂ | G | I/L | F |
| IBV | A | K | L | L | K | E | **R** | ✂ | G | F | F |
| ICV | V | T | K | P | K | S | **R** | ✂ | I | F | G |
| IDV | T | L | T | P | A | T | **R** | ✂ | I | F | G |

✂, protease cleavage site - arginine (R) indicated in bold typeface

To determine whether the proteases known to activate IAV HA also activated HEF, transduction titres of both IDV PVs produced by co-transfection of 293T cells with plasmids expressing HAT, TMPRSS2 or no protease were compared (Fig. 1A). For both IDV PVs produced with HAT, the transduction titres were higher than those with no protease, whereas the transduction titre obtained with PVs produced with TMPRSS2 were comparable to those of PVs produced with no protease or the delta envelope negative control. As all transduction titres were approximately 1 log higher with the HEF_Italy_ PV than with the HEF_France_ PV, further experiments were conducted with the HEF_Italy_ PV. The transduction titre of the HEF_Italy_ PV produced using exogenous trypsin at the plasmid transfection stage was also similar to that of the delta envelope negative control (Fig. 1B). Similarly, the transduction titres obtained with the ICV PV generated with trypsin or TMPRSS2 were low, comparable to the delta envelope negative control, whereas the titres obtained with PV produced with HAT were approximately 1.5-fold higher than the delta envelope negative control (Fig. 1C). Finally, the swine (sw) orthologues of human airway trypsin-like protease (swAT) and TMPRSS2 (swTMPRSS2) demonstrated similar proteolytic efficiency to their human orthologues in activating IDV PVs, with almost 2-fold higher transduction titres obtained for PVs produced with swAT than with sw-TMPRSS2 (Fig. 1D).

**Figure 1:**
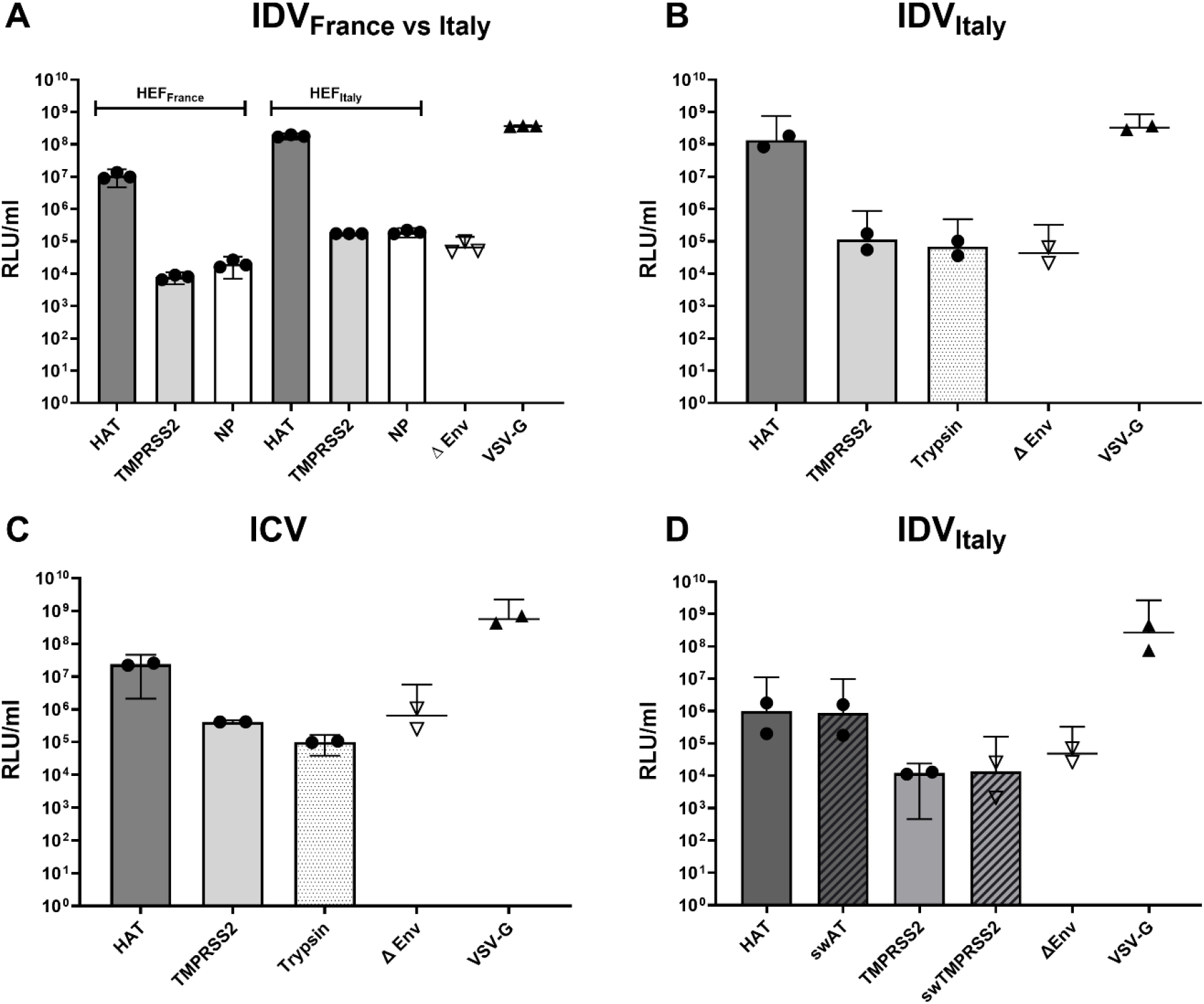
Comparison of activation of haemagglutinin-esterase fusion (HEF) proteins by proteases as measured by HEF-mediated pseudotyped virus (PV) transduction (A) influenza D virus HEF_France_ and HEF_Italy_ compared; (B) influenza D virus HEF_Italy_ (C) influenza C virus HEF; (D) influenza D virus HEF_Italy_. The PVs were produced using human or swine airway trypsin-like proteases (HAT and swAT) or TMPRSS2 (TMPRSS2 and swTMPRSS2) plasmids, exogenous trypsin or no protease (NP). Transduction titres expressed in mean relative light units per ml (RLU/ml) were determined 48 h after transduction of ST cells. PVs produced with no envelope gene (ΔENV) or VSV-G were used as negative and positive controls, respectively. In A, the results (mean and 95% CI of triplicate wells) of a representative experiment are shown. In B, C and D, the results (mean and 95% CI) of two biological replicates are shown.

When culturing infectious IDV, we observed that addition of exogenous trypsin was not required for multiple rounds of replication on MDCK cells whereas without trypsin, viral titres were 3 to 4 logs lower at 48-hours post-inoculation on A549 and MDBK cells, respectively (Fig. 2). A similar pattern was observed with A/PR8. This may be explained by the endogenous expression of matriptase by MDCK-II cells [29]. To confirm this, D/Italy/2015 PV was produced by co-transfecting ST cells with pcDNA3.1-matriptase. Although the transduction titre was lower for the PV produced using pcDNA3.1-matriptase compared with co-transfection with pCAGGS-HAT, it was higher than that of the PV produced with no protease (Fig. 3), suggesting that matriptase can also cleave the IDV HEF.

**Figure 2:**
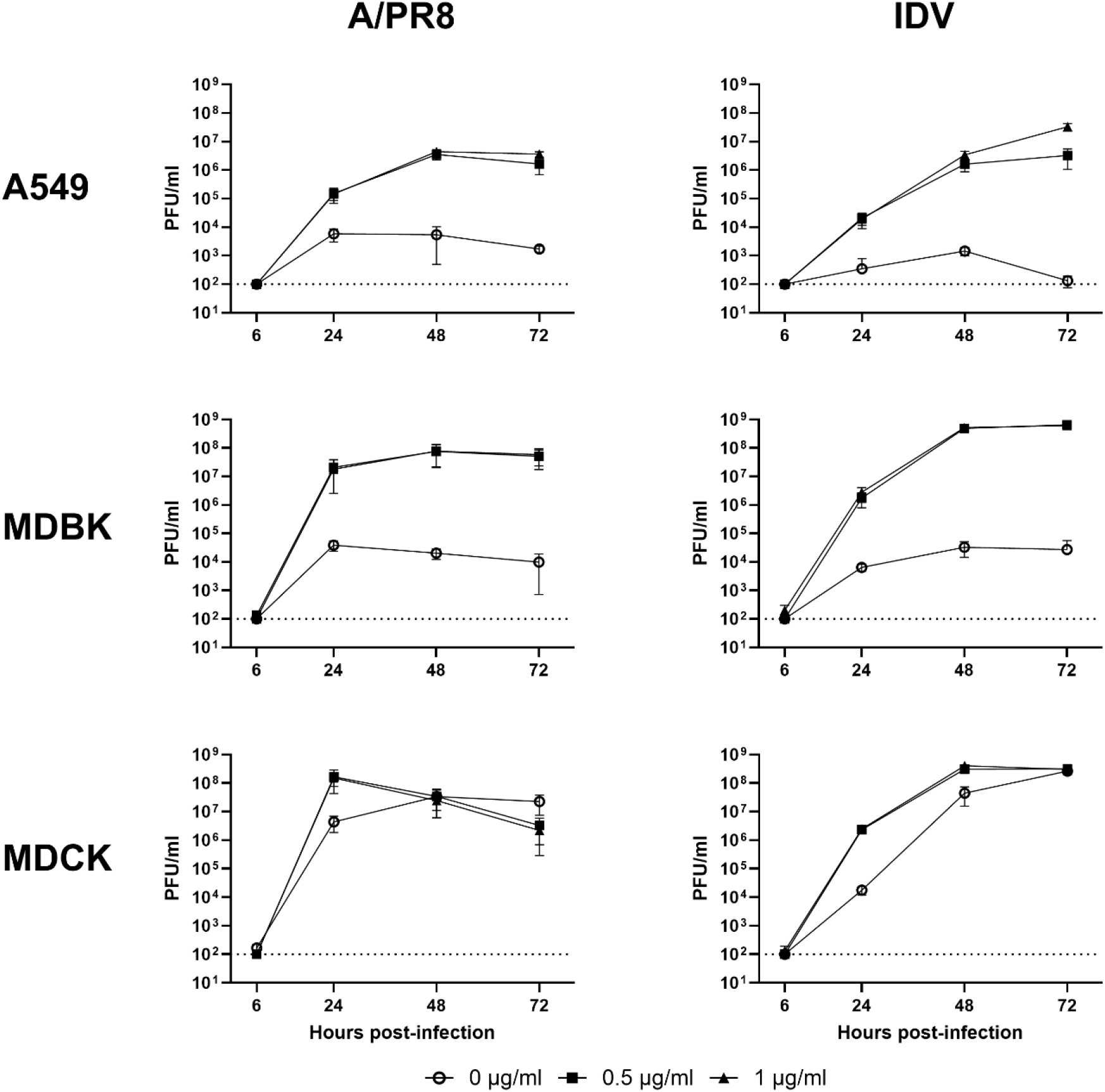
Multicycle replication of A/PR8 and influenza D virus on A549, MDBK and MDCK cells in the presence of varying quantities of TPCK-treated trypsin. The mean and SD of three biological replicates are shown. Titres are expressed as plaque forming units per ml (PFU/ml).

**Figure 3:**
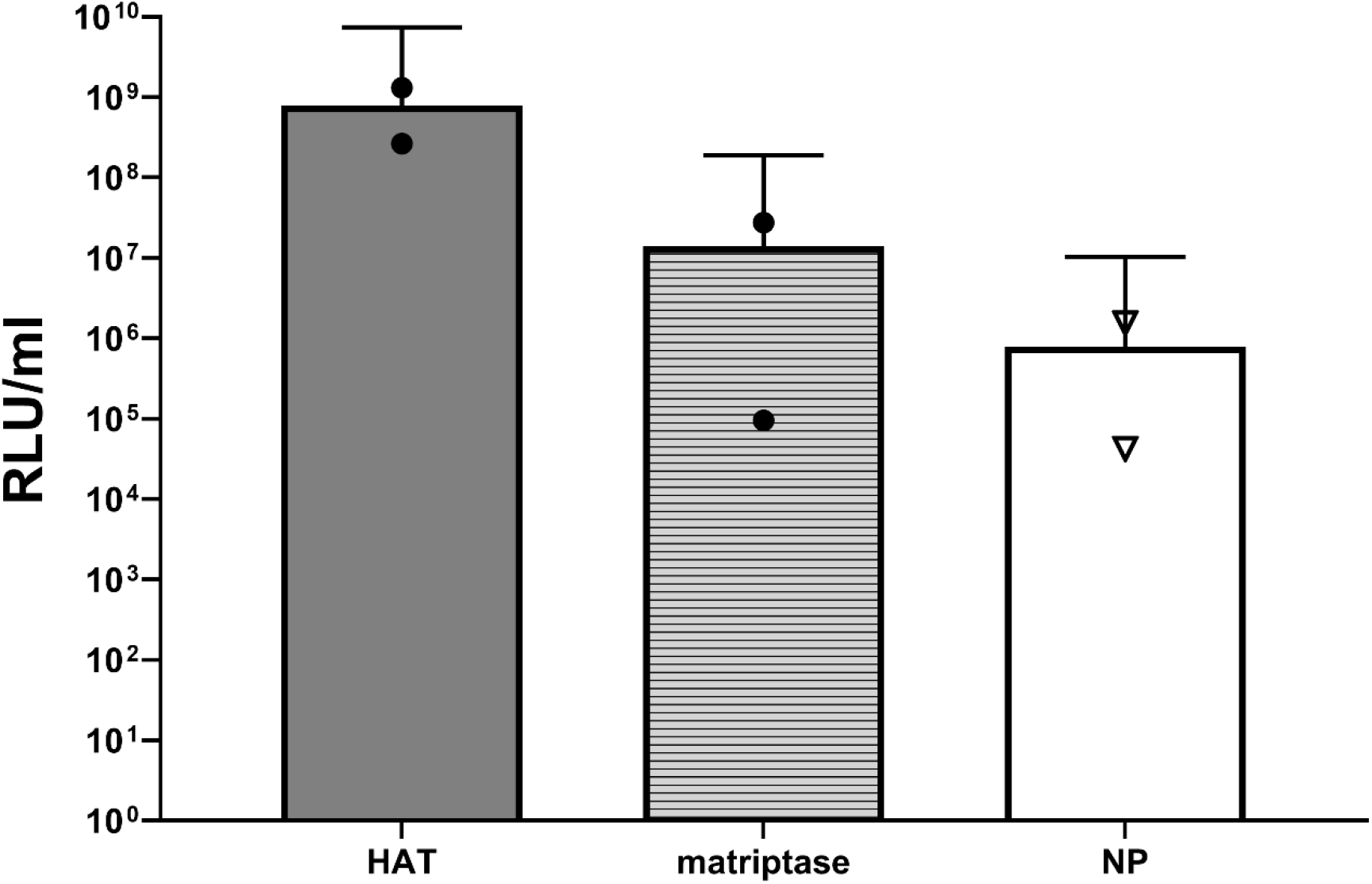
Activation of D/Italy/2015 pseudotyped virus by co-transfection with plasmid expressing matriptase. Pseudotyped virus generated without protease (NP) was used as negative control. Transduction titres expressed in relative light units per ml (RLU/ml) were determined 48 h after transduction of ST cells. the results (mean and 95% CI) of two biological replicates are shown.

## Discussion

The HEF proteins of ICV and IDV are synthesised as precursors that must be cleaved by cellular proteases. This cleavage enables fusion of the viral envelope with the endosomal membrane, which is responsible for viral infectivity and spread. The monobasic cleavage site of the HEF determines which protease cleaves the HEF, but which specific proteases that could cleave IDV HEF were hitherto unknown. Here, we confirmed, using HEF-bearing PVs, that the human airway trypsin-like protease (HAT) efficiently cleaves IDV HEF, whereas TMPRSS2 does not.

A notable finding of this study was that the swine analogue of HAT (swAT) activates IDV HEF with similar efficiency to HAT. Previously, Peitsch, Klenk [36] and Bertram, Heurich [32] showed that swAT and its chicken homologue could cleave IAV HA at monobasic cleavage sites with similar efficiency to the human protease HAT. Interestingly, the distribution and compartmentalisation of swAT and swTMPRSS2 in the porcine respiratory tract corresponds to that of HAT and TMPRSS2 in human respiratory tissues [36]. In addition, amino acid sequence alignment of human and swine airway proteases shows similarities of 81% (HAT/swAT) and 79% (TMPRSS2/swTMPRSS2). Bovine proteases are less well studied, particularly in the context of influenza viruses, but the amino acid similarities are like those for the swine orthologues: 75% for HAT and 83% for TMPRSS2.

As observed for IDV, the transduction titre of the ICV PV produced by co-transfection of pCAGGS-HAT was higher than when pCAGGS-TMPRSS2 was used to produce the PV. This contrasts with the only previous publication describing protease activation of ICV HEF [31]. In that study, MDCK cells were stably rather than transiently transfected with HAT or TMPRSS2 genes and the activation of ICV HEF in the context of a virus infection or PV transduction was higher in the TMPRSS2-expressing cells than in the HAT-expressing cells. However, the level of expression of HAT detected by western blot appeared to be substantially lower than of TMPRSS2 in the stably transfected cells. We could not confirm equivalent levels of expression of HAT and TMPRSS2 due to the lack of a tag, but we were able to compare expression levels of the swine counterparts as these were FLAG-tagged. This confirmed that the greater efficiency of HEF cleavage with swAT than with swTMPRSS2 was not due to higher expression – in fact the amount of protein expressed appeared to be higher for swTMPRSS2 than for swAT (data not shown). The use of the sequence of a different ICV isolate in our study and that of Sato et al. [31] could also contribute to the disparity. This is consistent with studies of IAV HA, in which the relative efficiency of cleavage by HAT or TMPRSS2 has been shown to vary among subtypes [39, 40].

Published studies [41, 42] support our observation that while exogenous trypsin is required for multiple cycles of replication of both A/PR8 and IDV in A549 and MDBK cells it is not required for MDCK cells. Godiksen, Selzer-Plon [43] have shown that MDCK cells endogenously express matriptase. Also, Hamilton, Gludish [41] have shown that matriptase cleaves the HA of H1N1 subtypes of IAV with the cleavage peptide sequence IPSIQSRGLF as shown in Table 1, including A/PR8. It is important to note that the MDCK strain used in these studies can affect findings. Baron, Tarnow [29] showed that MDCK-II cells, which are derived from higher passage MDCK cells and most commonly used in laboratories, can support replication of isolates of the H9N2 subtype, which have a dibasic cleavage site (R-S-S-R/R-S-R-R), whereas the earlier passage derived MDCK (H) strain, also known as MDCK I, cannot. This emphasises the importance of studying viral replication in different relevant cell lines.

Understanding the requirements for different proteases for viral activation can have potential clinical implications. Beaulieu, Gravel [44] demonstrated that a selective inhibitor of matriptase blocked IAV replication in human bronchial epithelial cells, suggesting that knowledge of which proteases activate influenza HA or HEF proteins could lead to the development of host-acting antiviral drugs.

In conclusion, we confirmed that the HEF of IDV must be proteolytically processed from its precursor form for a successful infection to occur and that it is efficiently cleaved by HAT. Although the IDV HEF was not apparently activated by TMPRSS2, we showed that matriptase can activate IDV HEF, to a lesser extent than HAT. Importantly, we demonstrated that cleavage of IDV HEF was similarly efficient whether using human or swine airway trypsin-like protease, suggesting that differences in protease cleavage of HEF is not a restriction factor in interspecies transmission of IDV.

## Acknowledgements

We thank Dr Mariette Ducatez (IHAP, Université de Toulouse, INRAE, ENVT, Toulouse, France) for providing D/bovine/France/5920/2014 and Dr Tom Peacock (The Pirbright Institute) for providing the pcDNA3.1-matriptase plasmid. We are grateful to Dr Stephen Dunham, Dr Kenneth Mellits, Dr Toshana Foster for support, helpful advice and discussions.

## References

1. ICTV. International Committee on Taxonomy of Viruses. https://ictv.global/taxonomy/ [accessed 10 June 2022 2021].

2. Francis Jr T, Quilligan Jr J, Minuse E. Identification of another epidemic respiratory disease. Science 1950;112(2913):495–495.

3. Taylor R. A further note on 1233 (“influenza C”) virus. Archiv für die gesamte Virusforschung 1951;4(4):485–485.

4. Kimura H, Abiko C, Peng G, Muraki Y, Sugawara K et al. Interspecies transmission of influenza C virus between humans and pigs. Virus research 1997;48(1):71–71.

5. Manuguerra JC, Hannoun C. Natural infection of dogs by influenza C virus. Research in virology 1992;143(3):199–199.

6. Zhang H, Porter E, Lohman M, Lu N, Peddireddi L et al. Influenza C virus in cattle with respiratory disease, United States, 2016–2018. Emerging infectious diseases 2018;24(10):1926.

7. Nissly RH, Zaman N, Ibrahim PAS, McDaniel K, Lim L et al. Influenza C and D viral load in cattle correlates with bovine respiratory disease (BRD): Emerging role of orthomyxoviruses in the pathogenesis of BRD. Virology 2020;551:10–15.

8. Matsuzaki Y, Katsushima N, Nagai Y, Shoji M, Itagaki T et al. Clinical features of influenza C virus infection in children. The Journal of infectious diseases 2006;193(9):1229–1229.

9. Salez N, Mélade J, Pascalis H, Aherfi S, Dellagi K et al. Influenza C virus high seroprevalence rates observed in 3 different population groups. Journal of Infection 2014;69(2):182–182.

10. Hause BM, Ducatez M, Collin EA, Ran Z, Liu R et al. Isolation of a novel swine influenza virus from Oklahoma in 2011 which is distantly related to human influenza C viruses. PLoS Pathog 2013;9(2):e1003176.

11. Oliva J, Mettier J, Sedano L, Delverdier M, Bourgès-Abella N et al. Murine Model for the Study of Influenza D Virus. Journal of Virology 2020;94(4):e01662–01619.

12. Hause BM, Collin EA, Liu R, Huang B, Sheng Z et al. Characterization of a novel influenza virus in cattle and Swine: proposal for a new genus in the Orthomyxoviridae family. mBio 2014;5(2):e00031–00014.

13. Su S, Fu X, Li G, Kerlin F, Veit M. Novel Influenza D virus: Epidemiology, pathology, evolution and biological characteristics. Virulence 2017;8(8):1580–1580.

14. Collin EA, Sheng Z, Lang Y, Ma W, Hause BM et al. Cocirculation of Two Distinct Genetic and Antigenic Lineages of Proposed Influenza D Virus in Cattle. Journal of Virology 2015;89(2):1036–1036.

15. Mekata H, Yamamoto M, Hamabe S, Tanaka H, Omatsu T et al. Molecular epidemiological survey and phylogenetic analysis of bovine influenza D virus in Japan. Transboundary and Emerging Diseases 2018;65(2):e355–e360.

16. Luo J, Ferguson L, Smith DR, Woolums AR, Epperson WB et al. Serological evidence for high prevalence of Influenza D Viruses in Cattle, Nebraska, United States, 2003-2004. Virology 2017;501:88–91.

17. Quast M, Sreenivasan C, Sexton G, Nedland H, Singrey A et al. Serological evidence for the presence of influenza D virus in small ruminants. Veterinary microbiology 2015;180(3-4):281–285.

18. Salem E, Cook EAJ, Lbacha HA, Oliva J, Awoume F et al. Serologic Evidence for Influenza C and D Virus among Ruminants and Camelids, Africa, 1991-2015. Emerg Infect Dis 2017;23(9):1556–1556.

19. White SK, Ma W, McDaniel CJ, Gray GC, Lednicky JA. Serologic evidence of exposure to influenza D virus among persons with occupational contact with cattle. Journal of clinical virology : the official publication of the Pan American Society for Clinical Virology 2016;81:31–33.

20. Nedland H, Wollman J, Sreenivasan C, Quast M, Singrey A et al. Serological evidence for the co-circulation of two lineages of influenza D viruses in equine populations of the Midwest United States. Zoonoses and public health 2018;65(1):e148–e154.

21. Shen M, Zhao X, Zhang J, Liu C, Qi C et al. Influenza D Virus in Domestic and Stray Cats, Northern China, 2024. Emerg Infect Dis 2025;31(8):1668–1668.

22. Trombetta CM, Fiori A, Falsini A, Pellegrini F, Le Poder S et al. Multicenter Serologic Investigation of Influenza D Virus in Cats and Dogs, Europe, 2015–2024. Emerging Infectious Disease journal 2026;32(2):293.

23. Mazzetto E, Bortolami A, Fusaro A, Mazzacan E, Maniero S et al. Replication of Influenza D Viruses of Bovine and Swine Origin in Ovine Respiratory Explants and Their Attachment to the Respiratory Tract of Bovine, Sheep, Goat, Horse, and Swine. Front Microbiol 2020;11:1136.

24. Nemanichvili N, Berends AJ, Wubbolts RW, Gröne A, Rijks JM et al. Tissue Microarrays to Visualize Influenza D Attachment to Host Receptors in the Respiratory Tract of Farm Animals. Viruses 2021;13(4).

25. Sreenivasan CC, Uprety T, Reedy SE, Temeeyasen G, Hause BM et al. Experimental Infection of Horses with Influenza D Virus. Viruses 2022;14(4):661.

26. Song H, Qi J, Khedri Z, Diaz S, Yu H et al. An Open Receptor-Binding Cavity of Hemagglutinin-Esterase-Fusion Glycoprotein from Newly-Identified Influenza D Virus: Basis for Its Broad Cell Tropism. PLoS Pathog 2016;12(1):e1005411.

27. Wang M, Veit M. Hemagglutinin-esterase-fusion (HEF) protein of influenza C virus. Protein Cell 2016;7(1):28–28.

28. Wang M, Ludwig K, Bottcher C, Veit M. The role of stearate attachment to the hemagglutinin-esterase-fusion glycoprotein HEF of influenza C virus. Cell Microbiol 2016;18(5):692–692.

29. Baron J, Tarnow C, Mayoli-Nüssle D, Schilling E, Meyer D et al. Matriptase, HAT, and TMPRSS2 activate the hemagglutinin of H9N2 influenza A viruses. J Virol 2013;87(3):1811–1811.

30. Bottcher E, Matrosovich T, Beyerle M, Klenk HD, Garten W et al. Proteolytic activation of influenza viruses by serine proteases TMPRSS2 and HAT from human airway epithelium. J Virol 2006;80(19):9896–9896.

31. Sato K, Hayashi H, Shimotai Y, Yamaya M, Hongo S et al. TMPRSS2 Activates Hemagglutinin-Esterase Glycoprotein of Influenza C Virus. J Virol 2021;95(21):e0129621.

32. Bertram S, Heurich A, Lavender H, Gierer S, Danisch S et al. Influenza and SARS-coronavirus activating proteases TMPRSS2 and HAT are expressed at multiple sites in human respiratory and gastrointestinal tracts. PloS one 2012;7(4):e35876.

33. Ferrara F, Molesti E, Böttcher-Friebertshäuser E, Cattoli G, Corti D et al. The human Transmembrane Protease Serine 2 is necessary for the production of Group 2 influenza A virus pseudotypes. Journal of molecular and genetic medicine: an international journal of biomedical research 2013;7:309.

34. Johnson MC, Lyddon TD, Suarez R, Salcedo B, LePique M et al. Optimized Pseudotyping Conditions for the SARS-COV-2 Spike Glycoprotein. J Virol 2020;94(21).

35. Marotta MG, Neto MM, Daly J, Maina M, van Diemen P et al. Development and optimisation of influenza C and influenza D pseudotyped viruses. J Virol Methods 2026;339:115243.

36. Peitsch C, Klenk HD, Garten W, Bottcher-Friebertshauser E. Activation of influenza A viruses by host proteases from swine airway epithelium. J Virol 2014;88(1):282–282.

37. Edgar RC. MUSCLE: multiple sequence alignment with high accuracy and high throughput. Nucleic acids research 2004;32(5):1792–1792.

38. Tamura K, Peterson D, Peterson N, Stecher G, Nei M et al. MEGA5: molecular evolutionary genetics analysis using maximum likelihood, evolutionary distance, and maximum parsimony methods. Molecular biology and evolution 2011;28(10):2731–2731.

39. Galloway SE, Reed ML, Russell CJ, Steinhauer DA. Influenza HA subtypes demonstrate divergent phenotypes for cleavage activation and pH of fusion: implications for host range and adaptation. PLoS pathogens 2013;9(2):e1003151.

40. Del Rosario JMM, da Costa KAS, Asbach B, Ferrara F, Ferrari M et al. Exploiting Pan Influenza A and Pan Influenza B Pseudotype Libraries for Efficient Vaccine Antigen Selection. Vaccines (Basel) 2021;9(7).

41. Hamilton BS, Gludish DWJ, Whittaker GR. Cleavage activation of the human-adapted influenza virus subtypes by matriptase reveals both subtype and strain specificities. Journal of virology 2012;86(19):10579–10579.

42. Godiksen S, Selzer-Plon J, Pedersen ED, Abell K, Rasmussen HB et al. Hepatocyte growth factor activator inhibitor-1 has a complex subcellular itinerary. Biochem J 2008;413(2):251–251.

43. Godiksen S, Selzer-Plon J, Pedersen EDK, Abell K, Rasmussen HB et al. Hepatocyte growth factor activator inhibitor-1 has a complex subcellular itinerary. Biochem J 2008;413(2):251–251.

44. Beaulieu A, Gravel É, Cloutier A, Marois I, Colombo É et al. Matriptase proteolytically activates influenza virus and promotes multicycle replication in the human airway epithelium. J Virol 2013;87(8):4237–4237.

